# Complex effects of pH on ROS from mitochondrial complex II driven complex I reverse electron transport challenge its role in tissue reperfusion injury

**DOI:** 10.1101/2020.08.31.275438

**Authors:** Alexander S. Milliken, Chaitanya A. Kulkarni, Paul S. Brookes

## Abstract

Generation of mitochondrial reactive oxygen species (ROS) is an important process in triggering cellular necrosis and tissue infarction during ischemia-reperfusion (IR) injury. Ischemia results in accumulation of the metabolite succinate. Rapid oxidation of this succinate by mitochondrial complex II (Cx-II) during reperfusion reduces the co-enzyme Q (Co-Q) pool, thereby driving electrons backward into complex-I (Cx-I), a process known as reverse electron transport (RET), which is thought to be a major source of ROS. During ischemia, enhanced glycolysis results in an acidic cellular pH at the onset of reperfusion. While the process of RET within Cx-I is known to be enhanced by a high mitochondrial trans-membrane ΔpH, the impact of pH itself on the integrated process of Cx-II to Cx-I RET has not been fully studied. Using isolated mitochondria under conditions which mimic the onset of reperfusion (i.e., high [ADP]). We show that mitochondrial respiration (state 2 and state 3) as well as isolated Cx-II activity are impaired at acidic pH, whereas the overall generation of ROS by Cx-II to Cx-I RET was insensitive to pH. Together these data indicate that the acceleration of Cx-I RET ROS by ΔpH appears to be cancelled out by the impact of pH on the source of electrons, i.e. Cx-II. Implications for the role of Cx-II to Cx-I RET derived ROS in IR injury are discussed.

## INTRODUCTION

Ischemia-reperfusion (IR) injury is caused by disruption of blood flow, followed by its restoration, and is relevant to pathologies such as myocardial infarction (heart attack) and stroke [1, 2]. The magnitude and duration of ischemia is a key determinant of the severity of eventual cell death, such that timely reperfusion is necessary to salvage tissue. However, paradoxically the reintroduction of oxygen by reperfusion triggers events including opening of the mitochondrial permeability transition (PT) pore, which initiate cell death [3].

Under physiologic conditions, pyruvate generated from glycolysis in the cytosol is shuttled into mitochondria and metabolized to acetyl-CoA to serve as a carbon source for the Krebs’ cycle. The cycle generates reducing intermediates NADH and succinate, that are consumed by complex I (Cx-I) and complex II (Cx-II) of the electron transport chain (ETC) respectively. These electrons then sequentially pass via Co-enzyme Q (Co-Q) to complex III (Cx-III), cytochrome *c*, complex IV (Cx-IV) and eventually O_2_. ETC activity pumps protons across the inner mitochondrial membrane (IMM), creating a protonmotive force (pmf), consisting of charge (ΔΨ_m_) and chemical (ΔpH) components, and ATP synthase (Cx-V) uses this pmf to produce ATP [4].

Several metabolic changes result from the lack of oxygen during ischemia, with the best-known being an elevation in glycolysis to yield lactic acid [5]. The release of a proton, yielding lactate, causes a drop in cytosolic pH from ~7.4 to ~6.6 during ischemia [6–8]. In addition, the accumulation of succinate is a hallmark of ischemia [9–11] and is proposed to occur via two mechanisms: (i) the reversal of mitochondrial Cx-II driven by a highly reduced Co-Q pool [9], and/or (ii) canonical Krebs’ cycle activity augmented by anaplerosis to α-ketoglutarate [12].

Within the first five minutes of reperfusion, accumulated succinate returns to normal levels, via a combination of its washout from tissue, and its rapid oxidation by Cx-II [12, 13]. The latter process leads to a highly reduced Co-Q pool, with the downstream ETC functioning at maximum capacity, resulting in an enhanced pmf [9]. These conditions provide the driving force to push electrons backward into Cx-I, a phenomenon known as reverse electron transport (RET) (Figure 1). [14–16] The generation of reactive oxygen species (ROS) has been observed under conditions that favor RET, and such ROS generation is thought to underlie the cascade of pathologic events including PT pore opening that lead to cell-death in IR injury [17]. Several variables have been identified that regulate RET, such as the redox states of Co-Q and NAD^+^/NADH pools, O_2_ concentration, ΔpH, and pH (for clarity, herein ΔpH refers to the trans-membrane proton gradient and pH refers to the pH in a given compartment) [18–21].

**Figure 1.**
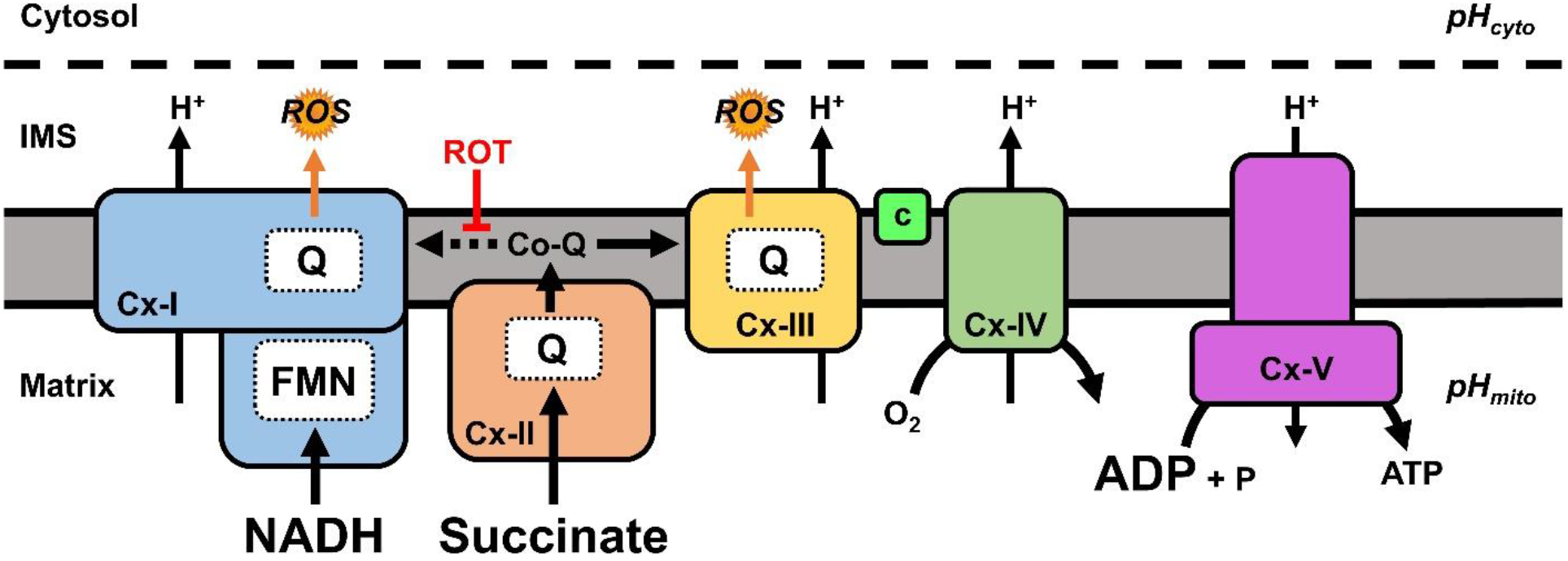
Schematic of mitochondrial ROS during reperfusion. At reperfusion, the re-introduction of O_2_ stimulates ETC activity and accumulated ischemic succinate is rapidly oxidized by Cx-II resulting in a highly reduced Co-Q pool. Given the metabolic conditions present at the start of reperfusion (high NADH and ADP), instead of flowing forward to Cx-III, some electrons can be driven backwards from Co-Q into Cx-I (reverse electron transport, RET, dotted line), generating ROS in a rotenone (ROT) sensitive manner. Abbreviations: pH_cyto_, cytosolic pH; pH_mito_, mitochondrial matrix pH; ETC, electron transport chain; Cx-I, complex-I; Cx-II, complex-II; Cx-III, complex-III; c, cytochrome c; Cx-IV, complex-IV; Cx-V, complex V; ROS, reactive oxygen species; IMS, intermembrane space; FMN, flavin mononucleotide; Co-Q, Co-enzyme Q; Q, Co-enzyme Q binding site.

The role of succinate oxidation by Cx-II as a driver of pathological ROS generation has been exploited for its potential as a drug target in IR injury [22, 23]. We and others have demonstrated that Cx-II inhibition at the start of reperfusion is protective in animal and perfused organ models of IR injury [12, 24–27]. In addition, it is known that prolongation of ischemic tissue acidosis into the reperfusion period can improve recovery from IR injury [28, 29]. While the exact protective mechanism of “acid post-conditioning” is poorly characterized, it may involve a recently described pH sensor on the PT pore [30]. Alternatively, the impact of pH on the major ROS generating sites within the ETC during reperfusion has not been extensively investigated.

RET-induced Cx-I ROS generation is thought to occur at the Co-Q binding site, and is characterized by sensitivity to Q-site inhibitors such as rotenone [18, 31–33]. The ΔpH is thought to be a major driver of RET-induced Cx-I ROS, although the effect of collapsing ΔpH on RET-induced Cx-I ROS is somewhat controversial [18, 20]. In addition, the relationship between cytosolic pH (pH_cyto_) and mitochondrial matrix pH (pH_mito_) and the independent effects of each on RET-induced Cx-I ROS are poorly understood. Importantly, many previous studies have assumed that the bulk of ROS measured under conditions of succinate supported respiration is due to Cx-I RET, without appropriate controls (i.e. subtraction of the rotenone-insensitive ROS component) [18, 20].

Herein, we examined the effects of pH_cyto_ range spanning ischemic to normoxic values (pH 6.6-7.8) on RET-induced Cx-I ROS (rotenone-sensitive), also taking into consideration the effects of pH_mito_ on the source of reduced Co-Q, i.e. Cx-II. Despite observing an inhibition of Cx-II at acidic pH, we observed no difference in the net production of RET-induced Cx-I ROS over the applied pH_cyto_ range. We conclude that while ΔpH may indeed serve to enhance RET-induced Cx-I ROS, this is counteracted by the impact of pH on the driving force (Cx-II). These findings have important implications for the impact of pH on ROS during reperfusion injury.

## RESULTS

### Acidic pH decreases mitochondrial succinate-linked respiration

To determine the effect of pH_cyto_ on mitochondrial respiration, we measured isolated mouse liver mitochondrial oxygen consumption rates (OCR) using a Seahorse XF96 analyzer over a pH range of 6.6 – 7.8. Mitochondria (5 μg protein / well) were assayed in the presence of 5 mM succinate to determine Cx-II linked respiration (state 2), followed by sequential injections of ADP (state 3) and rotenone/antimycin A (non-mitochondrial oxygen consumption). As the data in Figures 2B and 2C show, state 2 (quiescent) and state 3 (phosphorylating) respiration were lower at acidic pH_cyto_.

**Figure 2.**
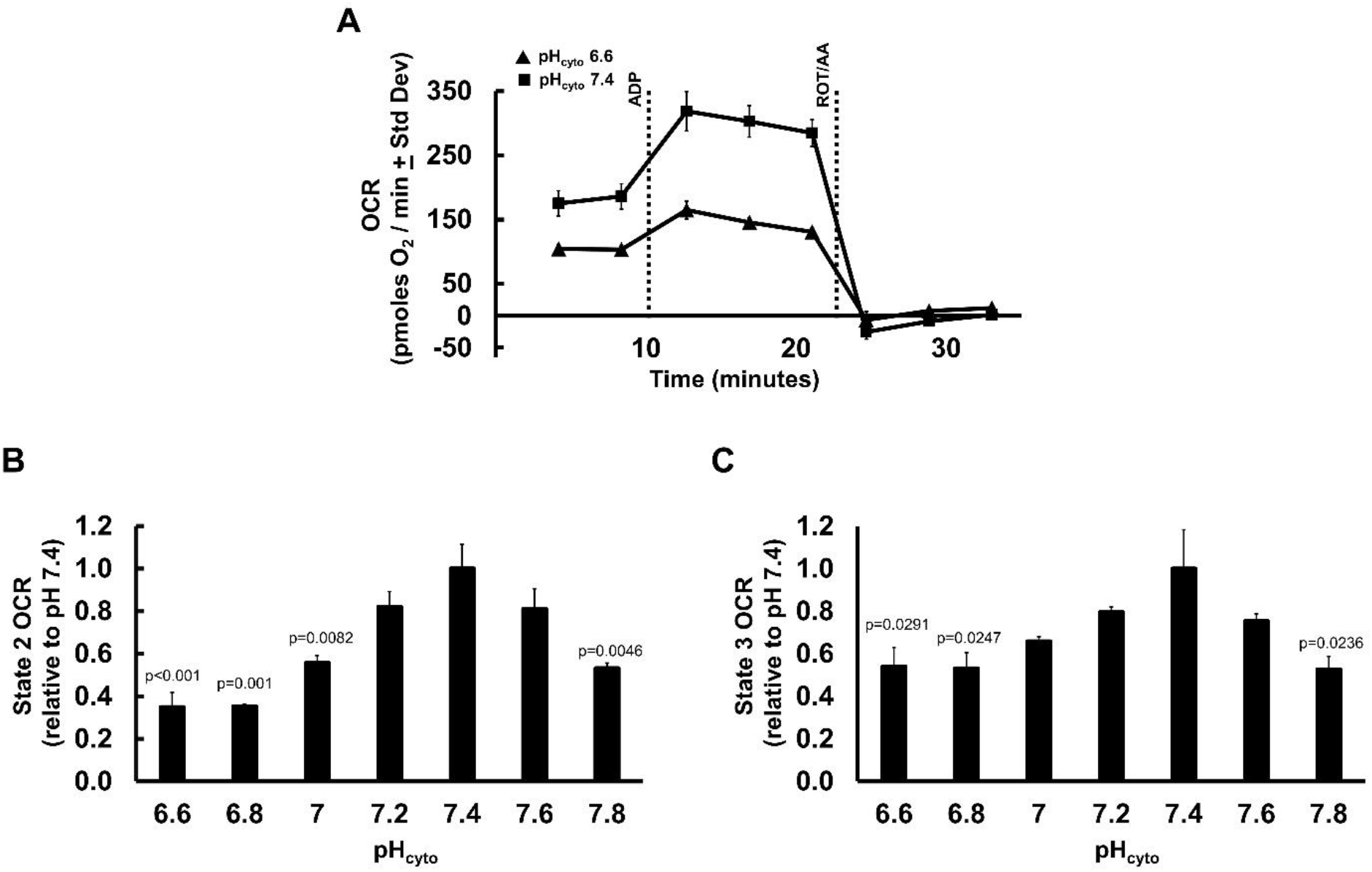
Mitochondrial respiration vs. pH_cyto_. (**A):** Representative Seahorse XF trace showing oxygen consumption rate (OCR) of mouse liver mitochondria in MRB pH_cyto_ 6.6 (triangles) and pH_cyto_ 7.4 (squares) sustained by succinate (state 2). ADP injection (state 3) and rotenone + antimycin A (ROT/AA) are indicated by dotted lines. Representative traces are means + Std. Dev. of one biological replicate with twelve technical replicates. **(B):** Oxygen consumption rates were assayed in isolated mouse liver mitochondria in succinate sustained state 2 respiration over pH_cyto_ range 6.6 – 7.8. **(C)**: Succinate + ADP sustained state 3 respiration over pH_cyto_ range 6.6 – 7.8. **(B and C)** Data are mean OCR values relative to individual OCR values at pH 7.4 for each mitochondria preparation + SEM (N = 4) with eight-twelve technical replicates per N. One-way ANOVA with Tukey’s test for multiple comparisons was performed. Statistical differences were determined between OCR values at all pH values, but values denoted in the figure are in comparison to pH 7.4.

### Acidic pH decreases complex II activity

Seeking the source of the observed acidic impairment of respiration, we next determined if acidic pH could impact the activity of mitochondrial Cx-II. Generally, the ΔpH across the IMM of mitochondria is 0.4–0.5 pH units, regardless of the prevailing pH_cyto_ [18, 20]. Thus, at normoxic or ischemic pH_cyto_ values of 7.4 or 6.6, the corresponding pH_mito_ values would be 7.8 and 7.0 respectively. Cx-II activity was assayed spectrophotometrically in freeze-thawed mouse liver or heart mitochondria over an applied pH range (i.e. pH_mito_) of 6.6–7.8 (equivalent to pH_cyto_ 6.2–7.4). As shown in Figures 3A and 3B, Cx-II activity was drastically reduced by acidic pH. These data suggest that under conditions of ischemic acidosis experienced during early reperfusion, Cx-II activity is low.

**Figure 3.**
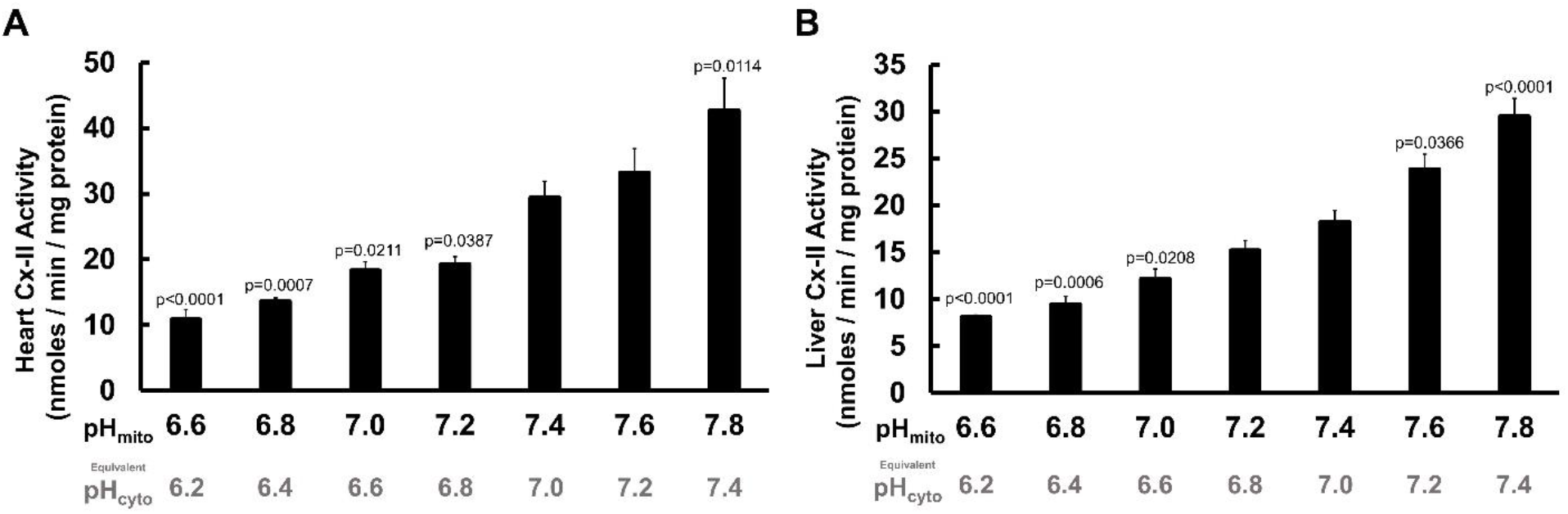
Isolated mitochondrial Cx-II activity vs. pH. Isolated complex-II (Cx-II) activity assays from freeze-thawed mitochondria demonstrated a pH-dependent relationship over the range pH 6.6-7.8. **(A):** Isolated Cx-II activity from heart mitochondria. **(B):** Isolated Cx-II activity from liver mitochondria. One-way ANOVA with Tukey’s test for multiple comparisons was performed. Data are means + SEM. In A, N = 6 biological replicates; four technical replicates per N. In B, N = 5 biological replicates; four technical replicates per N. Statistical differences were determined between all pH values, but values denoted in the figure are in comparison to pH 7.4. For each applied pH (i.e. pH_mito_), the equivalent pH_cyto_ is shown underneath, assuming ΔpH = 0.4 units.

### Acidic pH_cyto_ does not change net Cx-I RET ROS production

Incubation of mitochondria with succinate drives ROS generation both from Q site of Cx-I via RET, and from the downstream respiratory chain (likely the Q_O_ site of Cx-III). Since ROS generation from Cx-I RET can be inhibited by rotenone [18, 20], the contribution of Cx-I RET to the overall ROS signal is calculated as the rotenone-sensitive component. Metabolic conditions upon tissue reperfusion include high [succinate], high [NADH], and an elevated ADP/ATP ratio favoring state 3 respiration [34–36]. Initial experiments utilizing succinate in the presence of pyruvate plus carnitine to generate intra-mitochondrial NADH [12, 37] were unsuccessful owing to a quenching of the amplex red resorufin fluorescent signal by carnitine (data not shown). Further experiments using succinate in the presence of pyruvate plus malate for NADH production, yielded an elevation in ROS generation upon addition of rotenone, presumably due to enhancement of ROS production at the flavin site of Cx-I [38]. Thus, due to the confounding effect of NADH-generating substrates we opted to use succinate as the sole metabolic substrate, with ADP present to simulate early reperfusion.

Although a high ΔpH has been proposed as *sine qua non* for Cx-I RET [18], Figure 4A shows that even in the presence of ADP (state 3 respiration), ROS generation was partly rotenone-sensitive. Figure 4B shows the calculated net rates of rotenone-sensitive ROS (i.e., Cx-I RET) across the range of pH values studied, revealing that ROS from Cx-II driven Cx-I RET is not significantly enhanced at acidic pH.

**Figure 4.**
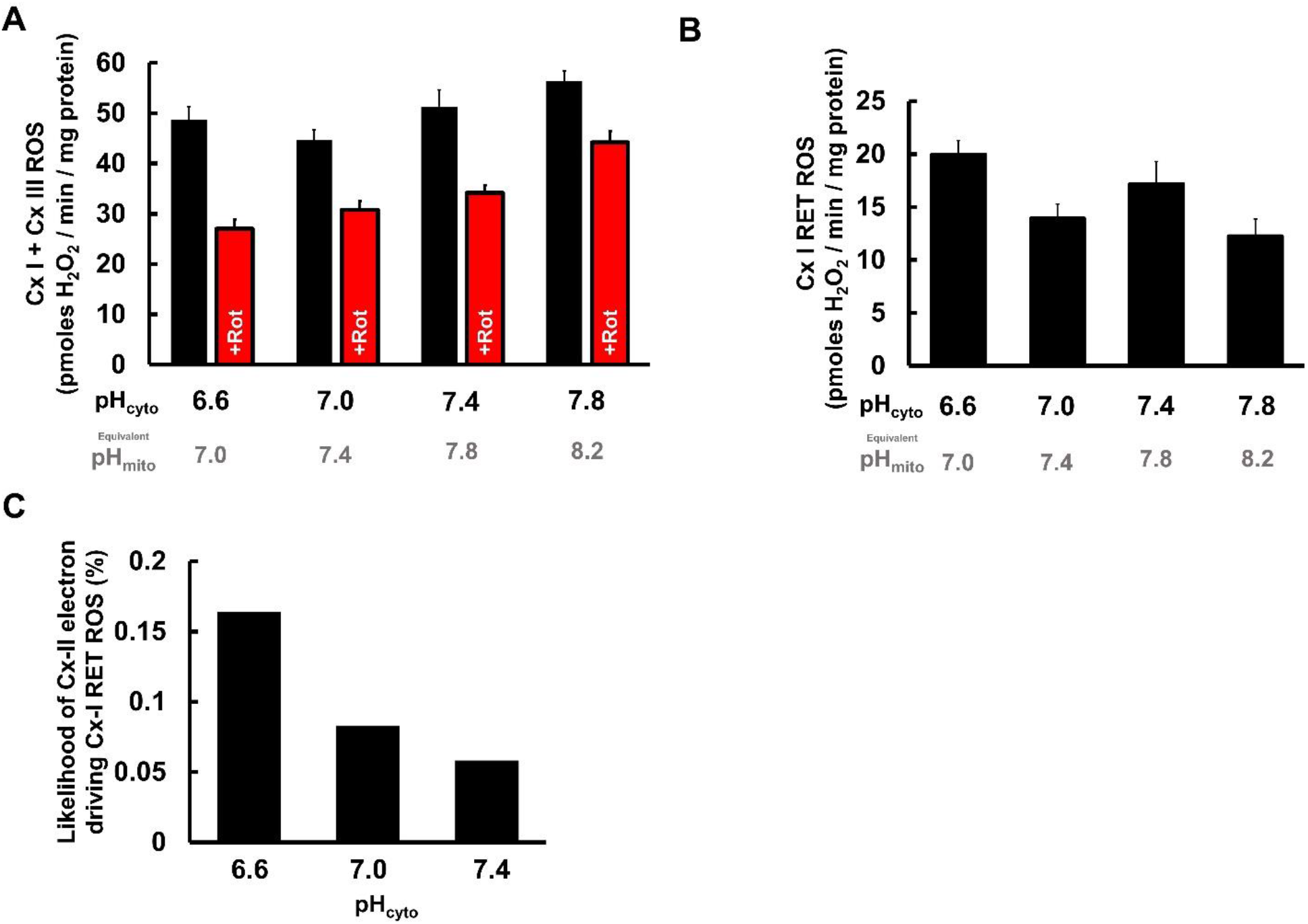
Cx-I RET ROS generation vs. pH_cyto_. **(A):** ROS production with or without rotenone (+Rot) in isolated liver mitochondria incubated with succinate and ADP across a pH range 6.6 – 7.8. **(B):** Net difference between ROS generated in the absence and presence of rotenone from panel B across the pH range. **(C):** Calculated probability that an electron oxidized from succinate at Cx-II (Fig. 3B) will go backward and cause Cx-I RET ROS (Fig 4B) across different values of pH_cyto_ 6.6 – 7.4. Data in A and B are means + SEMs (N=6) with six technical replicates per N. Data in C are calculated per Eqn. 1 (see text). For each applied pH (i.e. pH_cyto_), the equivalent pH_mito_ is shown underneath, assuming ΔpH = 0.4 units.

## DISCUSSION

The net amount of ROS originating from any site **(N)** is a combination of two factors: the availability of electrons **(E),** and the probability the site will leak an electron onto oxygen **(P)**.

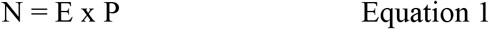

In the case of Cx-II driven Cx-I RET ROS, the availability of electrons in the Co-Q pool (E) is determined by the activity of Cx-II, and we measure the net ROS (N) directly from the rotenone-sensitive amplex red oxidation rate. So, the probability of Cx-I to donate electrons to molecular oxygen to make ROS (P) can be calculated by rearranging equation 1 (to P = N/E). As shown in Figure 4C, acidification of pH_cyto_ from pH 7.4 to 6.6 (equivalent pH_mito_ 7.8 to 7.0) results in a doubling of Cx-I RET ROS probability. This is in agreement with the widely demonstrated importance of ΔpH as a determinant of RET in Cx-I.

In the context of IR injury, the findings herein are particularly relevant to the phenomenon of acidic post-conditioning [28]. Specifically, our data suggest that the ability of acidic reperfusion to improve recovery from IR is likely not due to an inhibition of mitochondrial ROS generation. While Cx-II activity was inhibited at acidic pH, the probability that Cx-I can generate ROS via RET was enhanced, such that these two phenomena cancel each other out, yielding similar ROS generation rates across the range of pH values that would be expected in ischemia/normoxia.

A number of drugs developed to protect tissues such as the heart from IR injury are known to enhance cytosolic acidosis. For example, the cardioprotective drug cariporide inhibits the Na^+^/H^+^ exchanger (NHE1), an important pathway for export of protons from the cell [39, 40] Similarly, the NAD^+^ precursor NMN elicits cardioprotection in part via stimulation of glycolysis, which enhances lactic acidosis [41]. Furthermore, as discussed above, exogenously applied acidosis (i.e. acid reperfusion) is cardioprotective [28]. Contrasting the notion that Cx-I RET is enhanced by ΔpH, with the literature suggesting that acidic pH_cyto_ is cardioprotective, presents a paradox – how can acid be protective if it enhances RET driven ROS? However, factoring in the pH sensitivity of the source of electrons for RET during reperfusion, i.e., Cx-II, may explain these contrasting results: An acidic pH_cyto_ does not alter the overall Cx-II driven Cx-I RET derived ROS generation rate. This implies that the protective benefits of acidosis in the context of IR may originate elsewhere (e.g. effects on the PT pore) [30].

Regarding the mechanism by which acidic pH inhibits Cx-II itself, there is a possibility the protonation of succinate may render it an unsuitable substrate for the enzyme. However, succinate’s pKa values of 4.2 and 5.6 are such that at a physiologic pH_mito_ ~7.8, <1% of succinate is protonated as succinic acid, and even during ischemic acidification to pH_mito_ ~7.0, <5% of succinate would be protonated. Thus, succinate protonation is unlikely to account for Cx-II inhibition.

A surprising finding herein was that a significant portion of ROS generation with succinate as substrate was sensitive to rotenone, despite mitochondria respiring in state 3 (ADP present). Previously, in mitochondria from skeletal muscle, heart and brain, ~90% of the succinate-supported ROS generation was reported as rotenone-sensitive [18–20]. However, most of those studies used mitochondria in state 2 (no ADP), which does not model the situation at the beginning of reperfusion. As such, the observation herein that ~30% of the ROS signal is rotenone-sensitive and can thus be attributed to Cx-I RET, is likely a more accurate estimate of the contribution of this process to ROS generation during early reperfusion.

Recently, it was reported that application of an extracellular acidic pH can also impair the export of succinate from the cell via the monocarboxylate transporter MCT1[13]. It is unclear what may be the impact of the resulting succinate retention inside the cell, but it could be hypothesized that such retention would enhance IR injury due to succinate-driven ROS generation. However, the novel observation herein that acidic pH also inhibits Cx-II, may prevent retained succinate from being oxidized. Clearly, the complex dynamics of extracellular, intracellular, and mitochondrial pH in the first moments of reperfusion are poorly understood, and their further elucidation will be essential to determine the optimal manner in which succinate and Cx-II may be manipulated for therapeutic benefit in the context of IR injury.

## METHODS

### Animals and Reagents

Animal and experimental procedures complied with the National Institutes of Health *Guide for Care and Use of Laboratory Animals* (8^th^ edition, 2011) and were approved by the University of Rochester Committee on Animal Resources. Male and female C57BL/6J adult mice (8-12 weeks old) were housed in a pathogen-free vivarium with 12 hr. light-dark cycles and food and water *ad libitum*. Mice were administered terminal anesthesia via intra-peritoneal 2,2,2-tribromoethanol (Avertin) ~250 mg/kg.

### Heart Mitochondria isolation

Following anesthesia, the heart was excised and washed twice in 25 ml of ice-cold heart mitochondria isolation media (HMIM) comprising (in mM): Tris (20), EGTA (2), sucrose (300), pH 7.35. Tissue was diced using a razor blade, washed and transferred to a 50 ml conical tube containing 2 ml HMIM on ice, then homogenized using a Tekmar Tissumizer (IKA Instruments, Wilmington, NC) at 22,000 rpm for 20 s. The homogenate was centrifuged at 800 *g* for 5 min. at 4 °C, the pellet discarded, and the supernatant transferred to a new tube and centrifuged at 10,800 *g* for 5 min. Pellets were washed by centrifugation twice more, and the final pellet resuspended in 60 μl HMIM. Protein content was determined using the Folin-Phenol method against a standard curve constructed using bovine serum albumin [42, 43].

### Liver Mitochondria isolation

Following anesthesia, the liver was excised and washed in ice-cold 25 ml of liver mitochondria isolation media (LMIM) comprising (in mM): Tris (20), EGTA (2), sucrose (250), pH 7.35. The liver was chopped into 2-3 mm pieces using double scissors, washed 2-3 times and transferred into an ice-cold glass Dounce homogenizer. Tissue was first homogenized with 8-10 strokes using loose pestle A, then another 8-10 strokes using pestle B. Homogenates were centrifuged at 1,000 *g* for 3 min. at 4 °C the supernatant transferred to a new tube and spun at 10,000 *g* for 10 min. Pellet was resuspended in ice-cold LMIM and centrifuged at 10,000 *g* twice more and the final pellet resuspended in 600 μl of LMIM, transferred to a Potter-Elvehjem homogenizer and homogenized with 6-8 strokes. Protein content was determined using the Folin-Phenol method against a standard curve constructed using bovine serum albumin [42, 43].

### Isolated liver mitochondria respiration vs. pH

As previously described by Shiriai *et al*. [44] isolated liver mitochondria were loaded into a Seahorse XF96 microplate in 20 μl of mitochondria respiration buffer (MRB) pH 6.6 – 7.8 37°C (0.2 pH unit intervals). MRB contained the following (in mM): KCl (120), sucrose (25), MgCl_2_ (5), KH_2_PO_4_ (5), EGTA (1), HEPES (10), succinate (5). The loaded plate was centrifuged at 1000 *g* for 5 min at 4°C and an additional 130 μl of MRB (pH 6.6–7.8) was added to each well. For port injections, a 1 mM HEPES MRB pH 7.0 at 37°C was prepared, referred to as MRB_inj_. ADP (1.5 mM final) and rotenone/antimycin A (5 μM final) were prepared in 10x injection stocks in MRB_inj_. Mix and measure times were 0.5 and 3 min. Mitochondrial oxygen consumption rates (OCR) are expressed as pmols O_2_/min/mg mitochondrial protein.

### Complex II activity vs. pH

A Synergy 2 Multi-Detection 96-well Microplate Reader (BioTek, Winooski, VT) was used at 37 °C for experiments assessing complex II activity over the pH range 6.6–7.8. Isolated mitochondria (heart or liver) were freeze/thawed using liquid N2 three times. Mitochondria (0.1 mg/ml) were resuspended in 100 mM KPO4 assay buffer containing the following (in mM): Coenzyme-Q2 (0.05), EDTA (0.1), dichlorophenolindophenol (DCPIP) (0.12), KCN (1), rotenone (0.01) for 10 min at 37 °C in the plate reader. Activity was measured by spectrophotometrically monitoring the absorbance change of DCPIP at 600 nm (ε = 21 mM^−1^ cm^−1^). After baseline measurements, 5 mM succinate was added for reaction initiation and data collected every 50 s. for 5 min. Followed by addition of the Cx-II inhibitor 2-thenyltrifluoroacetone (TTFA, 1 mM) to determine the TTFA-sensitive rate [25].

### Measurement of ROS production by RET

A Synergy 2 Multi-Detection 96-well Microplate Reader (BioTek, Winooski, VT) was used at 37 °C for experiments assessing ROS generation as result of RET over a pH range. MRB (pH 6.6– 7.8) was supplemented with (in mM) Amplex Red (0.01), horse radish peroxidase (1 U/ml), superoxide dismutase (80 U/ml), ADP (0.1), succinate (2.5), ± rotenone (0.005), and incubated at 37 °C in the plate reader for 5 min. The reaction was initiated by addition of mitochondria (0.25 mg/ml), and ROS production measured as change in fluorescence from the conversion of amplex red to resorufin (λ_EX_ 570 nm, λ_EM_ 585 nm) when reacted with H_2_O_2_. To calibrate H_2_O_2_ production a known concentration of H_2_O_2_ was added at the end of each run. ROS generation was determined with or without the Cx I inhibitor rotenone, and the net rotenone-sensitive rate was used to represent Cx-I RET ROS [45, 46].

### Statistical Analysis

Comparisons between groups were made using ANOVA and with a post-hoc Tukey’s test applied where appropriate. Statistical analysis was performed using GraphPad Prism statistical analysis software. Data are shown as means ± SEM. Numbers of biological replicates (N) and technical replicates are noted in the figure legends. Significance was set at α = 0.05.

## Funding and additional information

This work was funded by a grant from the National Institute of Health (R01-HL071158). ASM was funded by an institutional NIH training grant (T32-GM068411). CAK was funded by a post-doctoral fellowship from the American Heart Association (#19POST34380212).

## Conflict of Interest

The authors declare that they have no conflicts of interest.

## ABBREVIATIONS

ROS: Reactive oxygen species
IR: Ischemia-reperfusion
mPTP: Mitochondrial permeability transition pore
ETC: Electron transport chain
Cx-I: Complex I (NADH:ubiquinone oxidoreductase)
Cx-II: Complex II (succinate:ubiquinone oxidoreductase, succinate dehydrogenase)
Cx-III: Complex III
Cx-IV: Complex IV
Cx-V: Complex V (ATP synthase)
Co-Q: Coenzyme Q (ubiquinone)
RET: Reverse electron transport
ΔpH: Transmembrane pH gradient
pmf: Proton motive force
ΔΨ_m_: Mitochondrial membrane potential
IMM: Inner mitochondrial membrane
IMS: Intermembrane space
pH_mito_: Mitochondrial matrix pH
pH_cyto_: Cytosolic pH
OCR: Oxygen consumption rate

## Notes

### Competing Interest Statement

The authors have declared no competing interest.

